# Untangling the role of selection and drift in population divergence via transcriptional network simulations: Extended analysis of Ghalambor et al. (2015)

**DOI:** 10.1101/277830

**Authors:** Kim L. Hoke, Kimberly A. Hughes, Eva K. Fischer, Cameron K. Ghalambor

## Introduction

A long-standing question in evolutionary biology is whether phenotypic plasticity influences adaptive evolutionary change (e.g. Waddington 1961; Price et al. 2003; Ghalambor et al. 2007; Lande 2009; Wund 2012). The same genotype can produce different phenotypes in response to different environments, but whether such plasticity constrains or facilitates evolutionary change remains an unresolved and controversial problem (Ghalambor et al. 2007). Theoretical models to date have made diverse predictions on the role of plasticity in evolutionary divergence (e.g. Ancel 2000; Paenke et al. 2007; Thibert-Plante and Hendry 2011), and empirical studies have largely been limited to retrospective approaches that infer the role of plasticity long-after populations and species have diverged (e.g. Losos et al. 2000; Wund et al. 2007; Scoville and Pfrender 2010). Field and lab studies that combine transcriptomic methods with recently diverged population comparisons provide a potentially powerful framework for quantifying patterns of plasticity for large numbers of molecular phenotypes within a generation, and how these phenotypes evolve across generations in response to either natural or artificial selection (e.g. Yampolsky et al. 2012; Ghalambor et al. 2015; Huang and Agrawal 2016). However, the high dimensionality of transcriptomic data imposes some computational challenges when attempting to infer the role of various evolutionary processes.

In Ghalambor et al. (2015) we reported rapid evolution of gene expression following colonization of a novel environment, and argued that non-adaptive plasticity resulted in stronger directional selection on a subset of genes. Specifically, we compared population differences in the brain transcriptomes of guppies (*Poecilia reticulata*) reared in a common garden one year after experimentally translocating guppies from streams with predators to replicate novel streams lacking predators. We focused on a subset of transcripts that were concordantly differentially expressed (CDE), in that they exhibited parallel divergence in the introduction populations and became more like a natural population lacking predators. We argued that these genes most likely evolved in response to selection, rather than by chance alone. We then examined the direction of plastic changes in this subset of genes when the source population was reared in the absence of predator cues and compared it to the direction of evolutionary change in the introduction populations. We found that the direction of plastic changes was largely in the opposite direction to the parallel evolved changes observed between the source and introduction populations. Such results suggested that genes exhibiting non-adaptive plasticity in a novel environment would be under stronger directional selection, whereas those exhibiting adaptive plasticity would be under weaker directional selection. In support of these conclusions, we designed custom permutation tests to generate null distributions against which to compare the observed data because chance fluctuations also would create negative relationships between plasticity and divergence. We based our conclusions on four lines of evidence or criteria: (1) the experimental data had a greater number of CDE transcripts than the permuted datasets; (2) the number of transcripts that diverged in opposite directions in the introduction populations (non-CDE) were not greater in the experimental data than in permuted datasets; (3) the correlation between divergence and plasticity was more negative in the experimental data than in permuted datasets; and (4) the number of CDE transcripts with plasticity and divergence in opposite directions was larger than in permuted datasets. Based on these criteria, we concluded the observed patterns were unlikely to have arisen by chance alone.

Two manuscripts have postulated alternative simple models that partially replicate these results without invoking a role for natural selection. If these results matched our experimental data, such models would cast doubt on our interpretation that natural selection produced the observed patterns. Specifically, Mallard et al. (2018) simulated transcriptional changes based on chance fluctuations, and van Gestel and Weissing (2018) posited a single regulatory change underlying population divergence of the set of CDE transcripts. Here, we expand on the analyses reported in Ghalambor et al (2018) in response to these manuscripts. Specifically, by exploring the details of these proposed alternatives, we argue that neither of these simple models replicates key features of the experimental data. In doing so, we bolster our original argument that natural selection against non-adaptive plasticity characterized the earliest stages of evolutionary divergence in the novel environments.

### Genetic drift model does not reproduce observed distributions

Mallard et al. (2018) claim their simulations can reproduce the results of Ghalambor et al (2015), however, as they acknowledge and we demonstrate, their claims fall short of meeting all four key criteria upon which the Ghalambor et al. (2015) results are based. Specifically, the simulations presented in the main text of Mallard et al (2018) do not account for criteria (1), and hence are irrelevant to consider. The simulations in their supplemental material require that parameter sets only meet criteria (1), (3), and (4) to be considered ‘significant’ – which forms the basis of their Supplemental Figure S3 – yet still do not consider criterion (2). We adapted their code to identify parameter sets that also meet criterion (2), (i.e. parameter sets that do not have more transcripts that diverged in opposite directions from each other in the two introduction populations (non-CDE transcripts) than in the permuted datasets). Many combinations of between-population and within-population variation yield simulated datasets in which both CDE and non-CDE transcripts exceeded the number in permuted datasets. Had our original dataset shown more non-CDE diverging transcripts than expected by chance, we would have concluded that random processes such as genetic drift had enabled extensive divergence in our introductory populations. Because only CDE genes were over-represented compared to the permuted datasets, we concluded that selection (both indirect and direct) was likely responsible for much of the divergence in our dataset. Ghalambor et al. (2018) report that only 26/400 parameter sets meet all four criteria, a small fraction of the parameter sets plotted in the comparable figure in Mallard et al (2018) that ignores essential criteria.

We next address Mallard et al.’s (2018) claim that the experimental data from Ghalambor et al. (2015) match their simulated distributions (Figure S4 in Mallard et al.). As above, we modified their code to represent the 26 parameter sets that met all four criteria. We compared two aspects of the distributions: (a) the total number of CDE genes (Figure 1A), and (b) the distribution of the between-population divergence to within-population divergence. In contrast to Mallard et al.’s (2018) assertion, very few parameter sets that meet all four criteria (Figure 1B, real data in green vs. simulated data in blue) have distributions consistent with the real data. In contrast, a large number of the parameter sets that do not meet all four criteria show substantial overlap (Figure 1c-real data in green vs. simulated data in blue).

**Figure 1:**
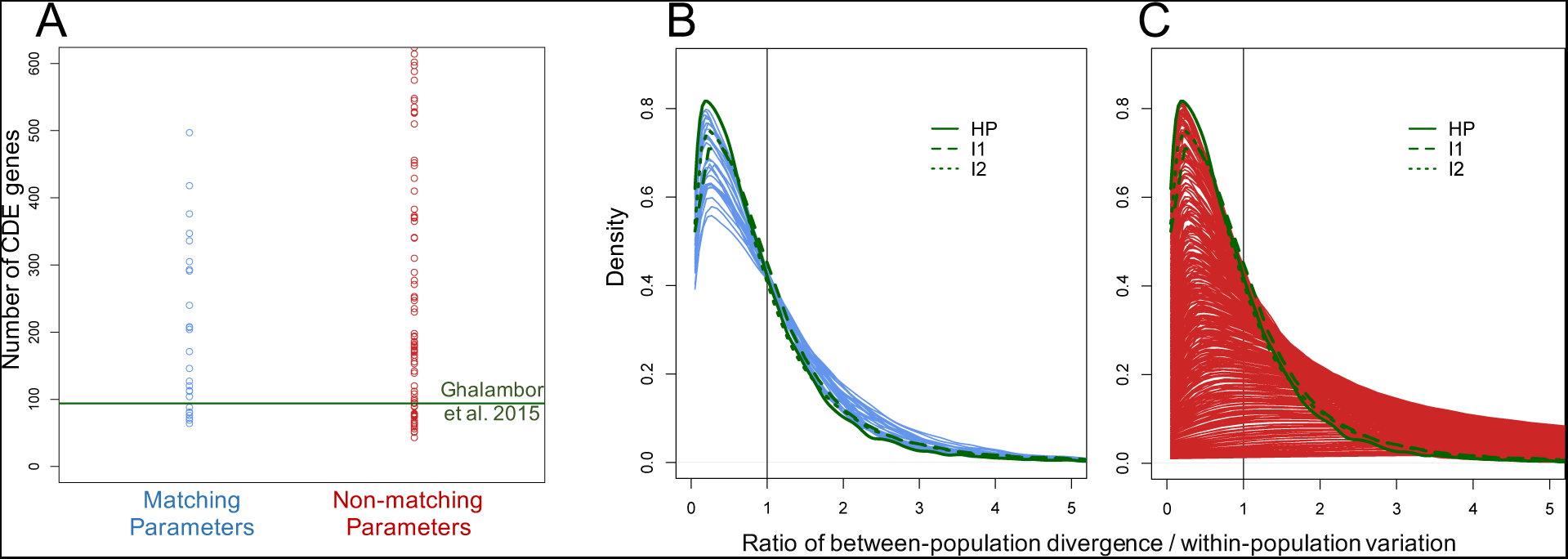
The distribution of the data from Ghalambor et al. (2015) matches only a fraction of the Mallard et al. (2018) simulations that meet all four criteria and matches a much larger number of parameter sets that do not replicate all four criteria. (A) Each dot represents the number of CDE genes predicted from simulations with a single set of parameters. Many of the simulations predicted much larger numbers of CDE transcripts than the 94 found in Ghalambor et al. (2015), marked by the green horizontal line, suggesting that the observed data do not come from a similar distribution. Not shown are the many simulations with non-matching parameters that predicted CDE numbers greater than 600. (B) We replot the distributions presented in Mallard et al. (2018) figure S4 to accurately depict the distributions of all simulations that met the four criteria (shown in blue) and (C) those that failed to meet at least one criterion (shown in red). Each curve represents the >30,000 genes from one simulation, summarizing the ratios calculated for each gene of the average group difference to the average variation around the mean within each group. Although all genes within any simulation share model-based average between- and within-population standard deviations, some genes will by chance have higher or lower values of this ratio, producing the spread in these curves. Note, the Mallard et al. (2018) figure appears very different because they plotted distributions from ¼ of the simulations, they overlaid red and blue curves, and they color-coded curves blue that failed criterion 2. The green curves represent observed values for the HP population and the two Introduction populations, as calculated by Mallard et al (2018). Note that a small subset of the 26 simulated curves plotted in panel B (simulations that met all four criteria, blue) have a reasonable match to the distribution of the original data (green). In contrast, a large number of curves that failed to recreate one or more criteria (panel C, red) overlap with the distributions from the original data (green).

In summary, only a tiny subset of the simulations outlined in Mallard et al. (2018) replicate the results from Ghalambor et al. (2015), but these cases are rare enough to be consistent with the permuted datasets that we used to draw all of our inferences. Thus, we conclude that the simulated datasets presented by Mallard et al. (2018) actually support, rather than refute, our conclusions that chance fluctuations are unlikely to have generated the observed patterns. For these reasons, we stand by our original analyses. We also strongly advocate for permutation-based approaches as in Ghalambor et al. (2015) rather than simulations for transcriptome analyses because permutations preserve the true distributions and covariances in the data.

### One regulatory change does not reproduce correlation structure observed in data

van Gestel and Weissing (2018) claim the results in Ghalambor et al. (2015) could be alternatively explained by a single potentially random regulatory change that mediated all the differential expression patterns in the CDE genes. To empirically test this assertion, we performed additional analyses to estimate the minimum number of changes and dimensionality within our data. If CDE genes diverged due a single regulatory change, we predicted all 94 CDE genes from Ghalambor et al. (2015) would be highly correlated with each other. Visualizing the correlations in our experimental data shows that many of our CDE transcripts are uncorrelated (Figure 2), although a few small sets of correlated genes are evident. We further examined correlations among all transcripts in our dataset (differentially expressed and not) using a weighted gene co-expression network analysis (Langfelder and Horvath 2008). This data-driven approach demarcated modules of highly correlated genes, and we assessed the distribution of CDE genes across these modules. If multiple regulatory changes were involved, we predicted CDE genes would be distributed across modules, whereas CDE genes would be confined to a single module if a single regulatory change caused the population differences. We found CDE genes were in fact distributed across modules, roughly in proportion to module size (Figure 3). Although these results do not identify the exact number of regulatory changes in these rapidly diverging populations, they do implicate multiple regulatory changes.

**Figure 2:**
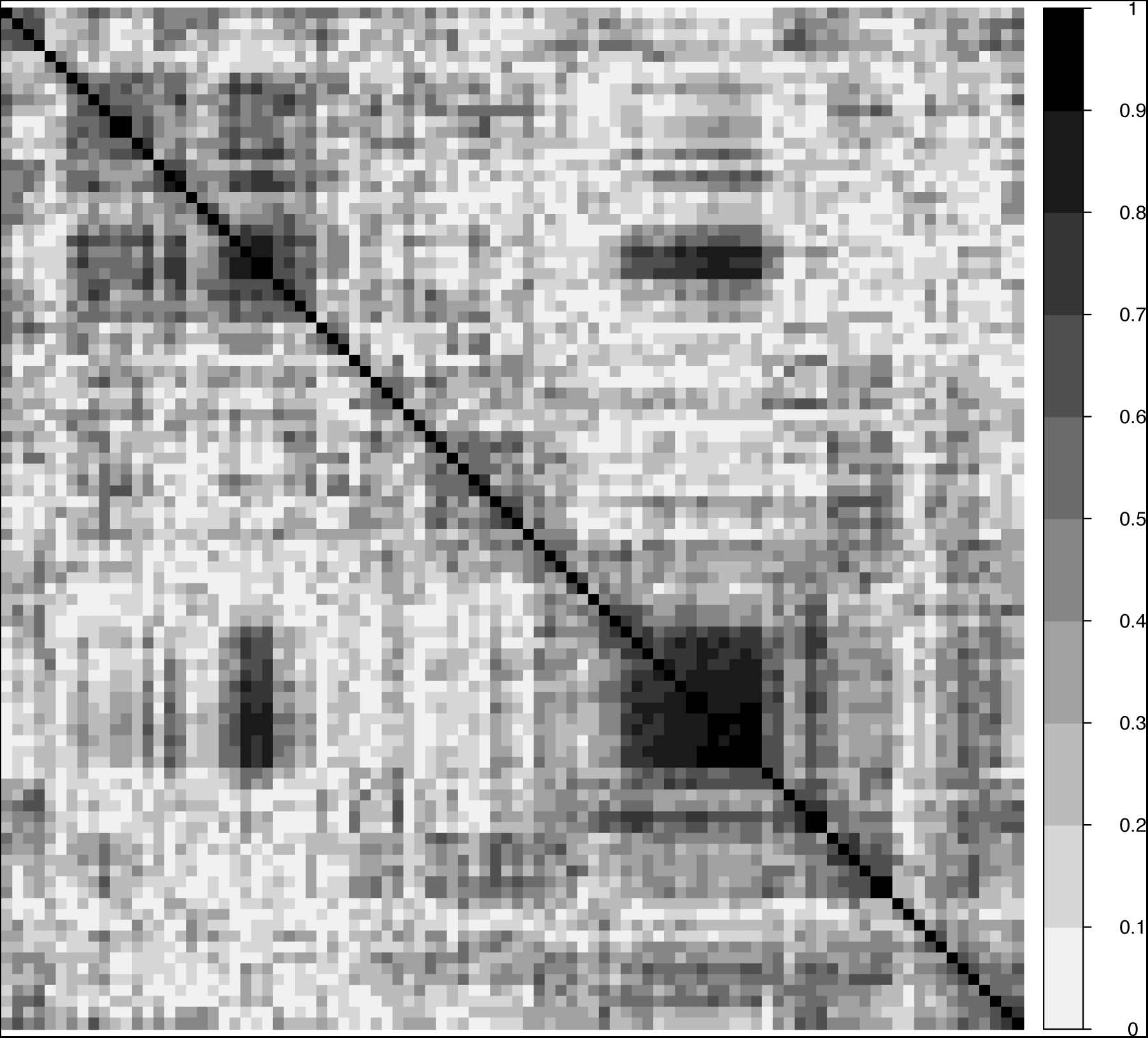
Correlation structure of CDE transcripts indicates numerous sets of strongly co-regulated transcripts. Each box represents a pairwise Pearson correlation, color-coded by strength as in the legend on the right. Transcript order was determined by hierarchical clustering to best visualize sets of likely co-regulated transcripts. Blocks of transcripts that are strongly correlated are visible along the diagonal. We suggest that these blocks likely share regulatory elements, and that some of these transcripts might have evolved due to indirect selection on correlated transcripts. The large number of very low correlations argue against a single causal modulator underlying divergence in all CDE transcripts.

**Figure 3:**
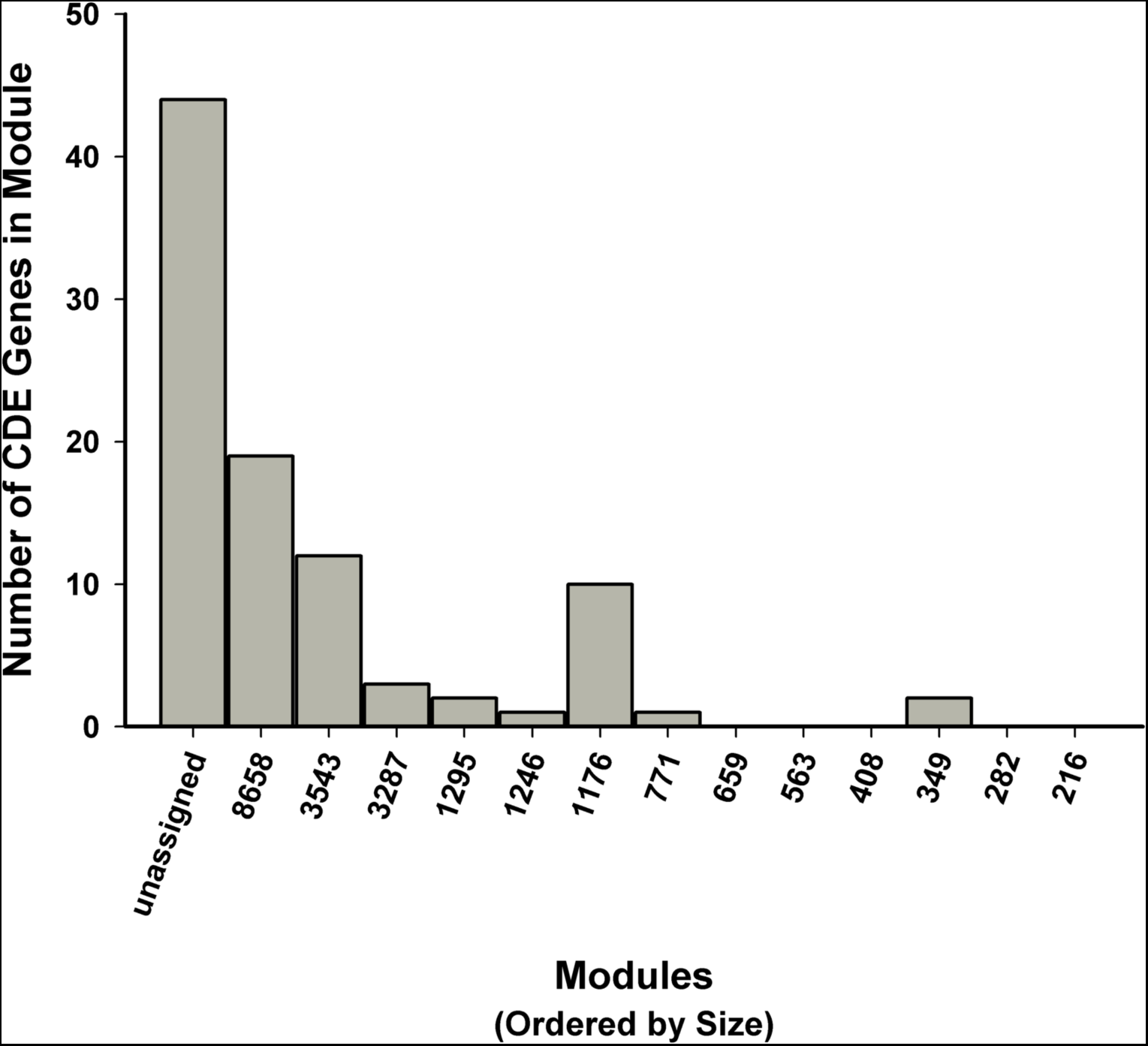
Frequency histogram showing WGCNA results. The thirteen modules calculated from the dataset using WGCNA are arranged by size (i.e. the total number of genes within each module, listed on the x-axis). The y-axis depicts the number of CDE genes within each module. Note that the CDE genes are spread across 8 well-defined modules, and nearly half the genes are not strongly associated with any module.

To further corroborate our findings, we used simulations to estimate the lowest pairwise correlation between genes consistent with van Gestel and Weissing’s (2018) proposal that CDE population differences were, in fact, due to a single shared modulator. We based our simulations on the proposed model (depicted in Figure 4) in which the mRNA levels of transcripts A and B both depend on the same hormonal regulator. In this model, all population differences in the transcripts are due to the population differences in the hormone. Importantly, these transcripts have other regulators (and noise) that do not vary among population; these other sources of regulation will decrease the pairwise correlations between transcripts. For our simulated datasets, we posited the parameter *w*, indicating how extensive the impact of the shared hormonal regulator is on abundance of transcripts, within the range 0.05 to 0.75, and varied the parameter *h* between 0.75 and 0.95 to indicate how closely hormone levels tracked group. We matched our experimental design as far as number of individuals in each group (population-rearing condition combination). We randomly selected group differences calculated from each of our CDE transcripts to set the mean for each group, multiplying the mRNA differences by factors ranging from 1 to 3 to account for decay in group differences due to noise and unexplained variance.

**Figure 4:**
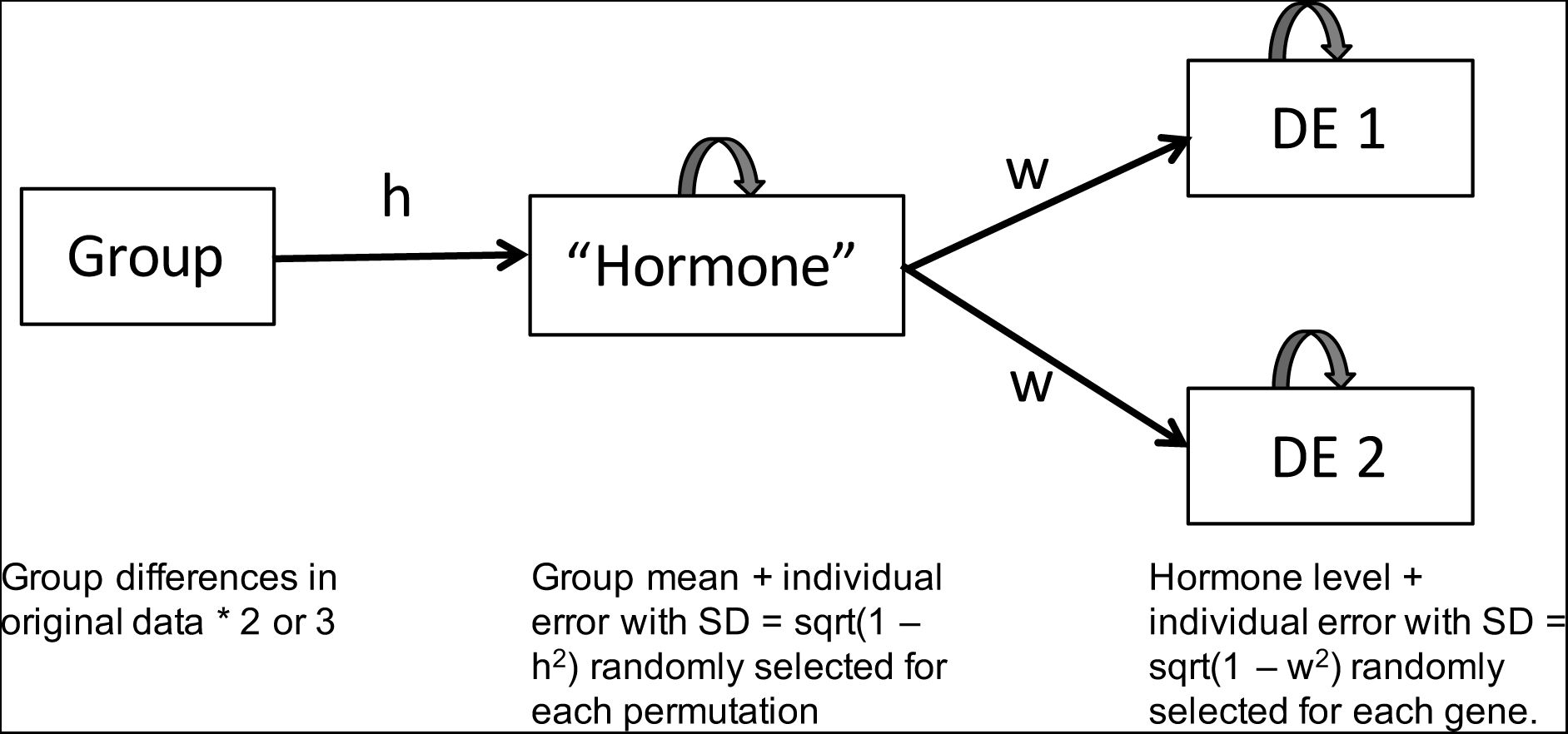
Simplified model used for simulating the relationship posited by van Gestel & Weissing (2018). The model postulates that group differences in all CDE genes (here, two are depicted, named DE1 and DE2, but simulations included up to 6000 transcripts) are due to shared reliance on an underlying regulator, here named ‘hormone’ for simplicity. The model predicts hormone levels in each of the 34 individuals in the dataset based on the group means from the original CDE transcripts for each population and rearing condition. We then estimate transcript abundance of all the dependent transcripts based on those hormone levels and transcript-specific noise (or unexplained variance due to other regulators that are not group-dependent). We determined which subset of the simulated transcripts would meet the differential expression criteria, then estimated the pairwise correlations among the simulated transcripts.

We sampled 600 (*w* ≥ 0.5) or 6000 (*w* < 0.5) transcripts that shared the same hormonal regulator from a normal distribution, with means determined by the model and standard deviations determined by sqrt(1-*h*^2^) for the hormone levels or sqrt(1-*w*^2^) for the transcripts (i.e. the unexplained variances for each parameter represented by the curved arrows in Fig. 4). We first assessed whether the simulated transcripts each showed concordant differential expression, accepting transcripts as CDE if t-tests for individuals from contrasting populations HP(source)- LP(natural low predation), HP(source)-Intro1(introduction 1), and HP(source)- Intro2(introduction 2) raised in non-predator conditions were each significant at p = 0.05 level. For the first 94 of those simulated transcripts that met the CDE criteria, we calculated the pairwise Pearson correlation matrix of all transcripts across all individuals. The results of these simulations are shown in Figure 5, and highlight the discrepancy between the results of the simulations with the observed distribution of correlation strength (i.e. the absolute values of the pairwise correlations among CDE transcripts-shown as a green line in Figure 5).

**Figure 5:**
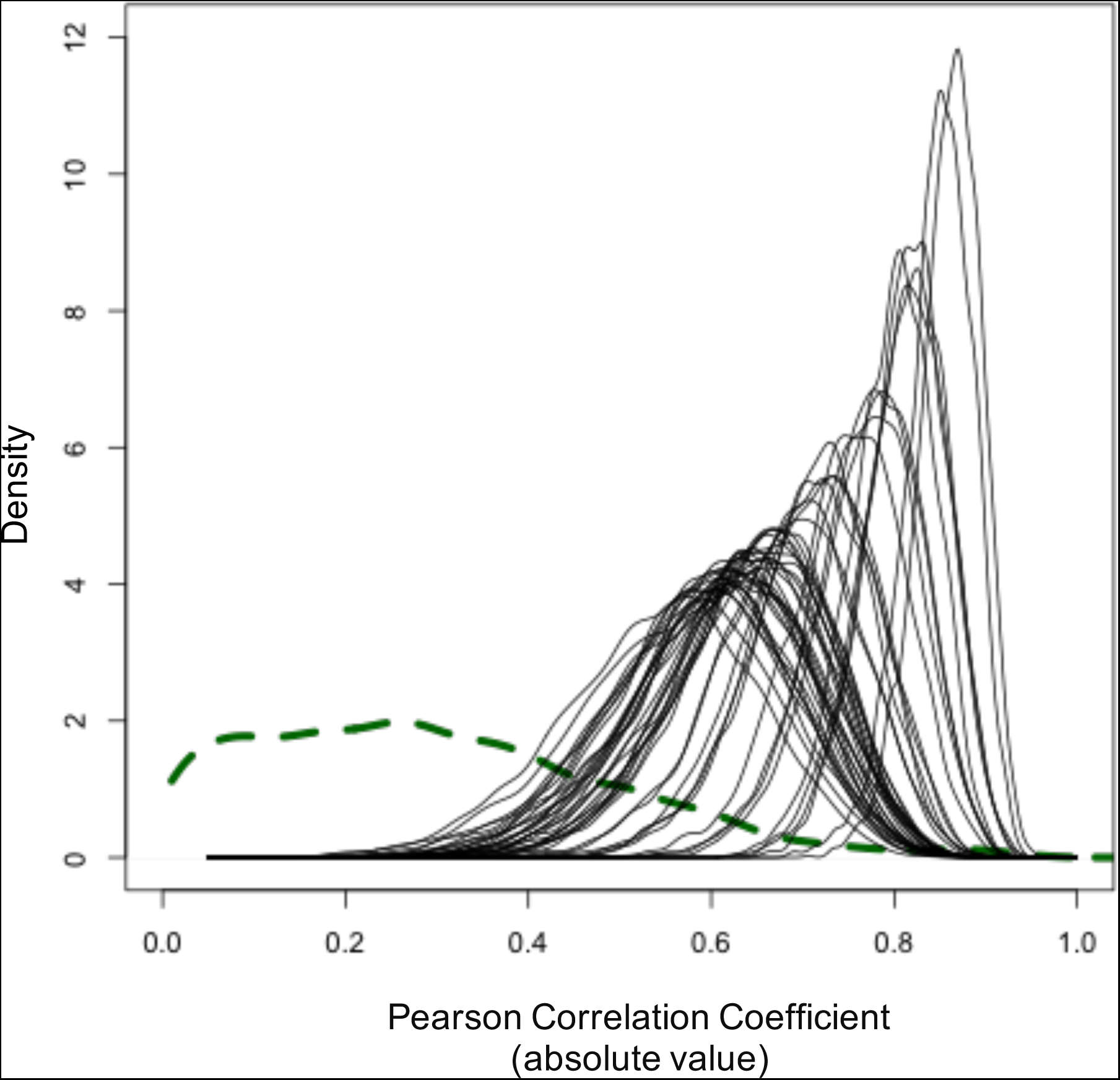
Results of simulations from a wide range of parameters demonstrate substantially higher distributions of correlations than are found in the original data. Black curves represent the distribution of pairwise correlations in simulated CDEs derived from a single parameter combination (only simulations with at least 94 CDE genes are included). Even for parameter values positing very low correlations between the hormone levels and dependent transcripts, nearly all simulated genes that met the criteria to be called CDE had correlation values above 0.4. In contrast, the green curve represents the distribution of correlations among CDE genes in the original dataset, a distribution that does not match any of the simulated data.

In summary, our complementary correlation analyses demonstrate that numerous regulatory changes give rise to the 94 CDE genes in Ghalambor et al. (2015). Our data are not consistent with the proposal by van Gestel & Weissing (2018) that one or a very small number of changes underlie the broad patterns in the dataset. Thus, in combination with statistical tests comparing the divergence in gene expression compared to putative neutral markers (see Extended Data Table 1 in Ghalambor et al. 2015), strong evidence suggests that selection, rather than random processes like drift, were responsible for generating rapid and adaptive changes in gene expression.

